# Discovery of potential imaging and therapeutic targets for severe inflammation in COVID-19 patients

**DOI:** 10.1101/2020.07.20.213082

**Authors:** Hyunjong Lee, Hyung-Jun Im, Kwon Joong Na, Hongyoon Choi

**Author notes:** [Correspondence and Reprint Request] Hongyoon Choi, MD., PhD., Department of Nuclear Medicine, Seoul National University Hospital, 101 Daehak-ro, Jongno-gu, Seoul, Republic of Korea, 03080, Kwon Joong Na, MD., Department of Thoracic and Cardiovascular Surgery, Seoul National University Hospital, 101 Daehak-ro, Jongno-gu, Seoul, Republic of Korea, 03080, Hyung-Jun Im, MD, PhD, Department of Transdisciplinary Studies, Graduate School of Convergence Science and Technology, Seoul National University, Gwanggyo-ro 145, Yeongtong-gu, Suwon-si, Gyeonggi-do, Republic of Korea, 16229.

## Abstract

The COVID-19 pandemic has caused more than 540,000 deaths globally. Hyperinflammation mediated by dysregulated monocyte/macrophage function is considered to be the key factor that triggers severe illness in COVID-19. However, no specific targeting molecule has been identified for detecting or treating hyperinflammation related to dysregulated macrophages in severe COVID-19. Herein, we suggest candidate targets for imaging and therapy in severe COVID-19 by analyzing single-cell RNA-sequencing data based on bronchoalveolar lavage fluid of COVID-19 patients. We found that expression of *SLC2A3*, which can be imaged by [^18^F]fluorodeoxyglucose, was higher in macrophages from severe COVID-19 patients. Furthermore, by integrating the surface target database and drug-target binding database with RNA-sequencing data of severe COVID-19, we identified *CCR1* and *FPR1* as surface and druggable targets for drug delivery as well as molecular imaging. Our results provide a resource for candidate targets in the development of specific imaging and therapy for COVID-19-related hyperinflammation.

## Introduction

The Coronavirus disease 2019 (COVID-19) pandemic caused by severe acute respiratory syndrome coronavirus 2 (SARS-CoV-2) infection has led to over 540,000 deaths worldwide as of July 9, 2020 (*1*) (https://coronavirus.jhu.edu/map.html). The Chinese Centers for Disease Control and Prevention analyzed the characteristics of 72,314 cases of COVID-19 and reported that disease severity varies widely, with 81% of patients experiencing mild disease, 14% developing severe disease, and 5% developing critical disease that is characterized by respiratory and/or multiorgan failure (*2, 3*). In most cases, severe/critical disease develops within 2 weeks after symptom onset (*4*), and in a recent study, it was reported that the mortality of patients undergoing mechanical ventilation was 88.1% (*4, 5*). Therefore, efforts to identify and manage patients who are at high risk of developing severe illness are urgently needed.

Hyperinflammation mediated by dysregulated macrophage and monocyte responses is considered to be the key factor causing severe disease in patients with severe COVID-19 based on observations of prevalent lymphocytopenia and massive infiltration of macrophages in multiple organs, including the lungs, spleen, lymph nodes, heart and kidney (*3, 6*). Recently, targeted probes based on small molecules and antibodies have contributed to precise diagnosis and treatment in various inflammatory and infectious diseases (*7, 8*). Nonetheless, which molecular targets might be utilized in imaging and drug delivery for hyperinflammation in severe COVID-19 has not been investigated. Molecular imaging to detect characteristic hyperinflammation by a dysregulated immune response has the potential to predict the progression of severe COVID-19. Furthermore, a drug delivery system to target specific immune cells that cause hyperinflammation in severe COVID-19 might allow precise immune modulation and greatly impact patient survival.

In this study, we analyzed single-cell RNA-sequencing (scRNA-seq) data based on bronchoalveolar lavage (BAL) fluid cells of healthy controls and COVID-19 patients along with three different databases, the Surfaceome database (*9*), the Database of Imaging Radiolabeled Compounds (DIRAC) (*10*), and BindingDB (*11*), to identify feasible targets for molecular imaging and therapy in severe COVID-19.

## Results

### Composition of immune cells in BAL fluid

scRNA-seq data of BAL fluid cells were obtained from three patients with moderate COVID-19, six patients with severe/critical COVID-19, and four healthy controls (Gene Expression Omnibus database, accession number GSE145926) (*12*). BAL fluid cells were classified into 21 clusters according to expression of cell-type specific marker genes (Fig. 1A, Supplementary Fig. 1). We explored the cell subpopulation of each group and identified a substantial difference in the composition of immune cells between the three groups (Fig. 1B). The majority of cells in BAL fluid were composed of different types of macrophages in the three groups. T-cells were more abundant in the moderate COVID-19 group than in the severe COVID-19 group. In contrast, neutrophils, natural killer (NK) T-cells, and plasma cells were more abundant in the severe COVID-19 group than in the moderate COVID-19 group. The composition of macrophage subpopulations differed between the three groups: 1) in the healthy control group, a cluster of macrophages, M02, was the most abundant subpopulation; 2) in moderate COVID-19, M04 was the most abundant cluster; and 3) in severe COVID-19, M01 and M03 were the two most dominant cell types.

**Fig. 1.**
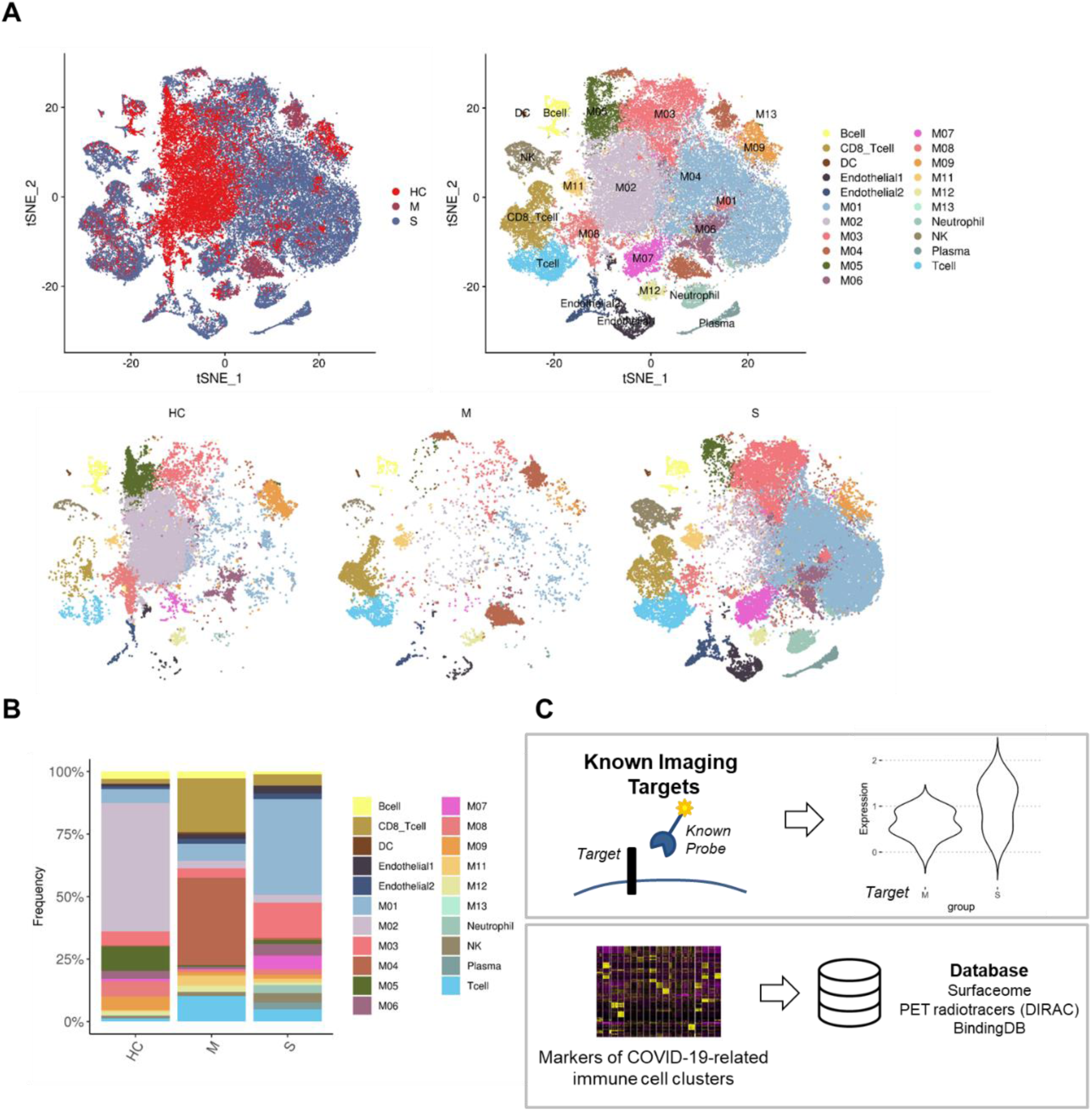
Bronchoalveolar immune landscape of COVID-19 patients and overview of the study design. **(A)** t-SNE projection of major cell type clusters in BAL fluid according to the severity of COVID-19 infection. Each point represents a single cell, and color coding of each patient group (*upper left*) and cell type population (*upper right*) are shown adjacent (HC = healthy control; M = moderate COVID-19 infection; S = severe COVID-19 infection). **(B)** The composition of major cell types per patient group. **(C)** Workflow for targetable molecule discovery in severe COVID-19 patients. We analyzed the expression level of potential imaging targets for COVID-19 patients. Additionally, we explored the marker genes of COVID-19-related immune cell clusters from three public databases (Surfaceome database, PET tracer database, and BindingDB).

We utilized the following two different approaches to discover potential targets to image and/or modulate the dysregulated immune system in severe COVID-19: 1) evaluating the expression levels of alleged imaging targets for inflammation and 2) exploring markers filtered by targetable molecules based on publicly available databases (Fig. 1C).

### Evaluating expression levels of alleged imaging targets

We evaluated the expression levels of the molecular targets of clinically available imaging tracers for inflammation in BAL fluid cells of COVID-19. First, we selected molecular targets and matched imaging tracers among metabolism-related tracers: glucose transporters (GLUT)/2-[^18^F]-fluoro-2-deoxy-d-glucose ([^18^F]FDG) (*13*), monocarboxylate transporters (MCT)/[^11^C]-acetate (*14*), folate receptors (FOLR)/[^18^F]-labeled folic acid derivatives (*15*), and L-type amino acid transporter 1/[^11^C]-methionine (*16*). Furthermore, target molecules for macrophage imaging and matched imaging tracers were selected: translocator protein (TSPO)/[^11^C]-PBR28 (*17*) and mannose receptor 1 (MRC1)/2-deoxy-2-[^18^F]-fluoro-D-mannose (*18*). The expression levels of other imaging target molecules were also examined: somatostatin receptor subtype-2 (SSTR2)/[^68^Ga]-DOTA-TATE (*19*), fibroblast-activated protein (FAP)/[^68^Ga]-fibroblast-associated protein inhibitor (*20*) considering pulmonary fibrosis of COVID-19 (*21*), alpha-v-beta-3 integrin (ITGAV)/arginylglycylaspartic acid (RGD) (*22*), CD8+ T-cells/[^89^Zr]-radiolabeled human CD8-specific minibody (*23*), and granzyme B (GZMB)/[^68^Ga]-NOTA-GZP (*24*).

All selected molecular targets were projected onto a t-distributed stochastic neighbor embedding (t-SNE) plot depicting 21 immune cell clusters of all patients, and macrophages enriched in severe COVID-19 patients were marked separately (Fig. 2A). The expression levels of molecular targets were scattered separately for cells from healthy controls, moderate COVID-19, and severe COVID-19 for each immune cell cluster. *SLC2A1* (GLUT1) showed low expression across the immune cell clusters, whereas *SLC16A3* (MCT4) was highly expressed in macrophage clusters. To analyze whether molecular targets are able to distinguish the severity of COVID-19 infection, we compared the expression level of each target between the groups (Fig. 2B, Supplementary Fig. 2). We found that *SLC2A3* (GLUT3) was increased in macrophage clusters enriched in severe COVID-19 patients. TSPO and MRC1 were highly expressed in macrophage clusters enriched in healthy controls and moderate COVID-19 patients. GZMB was highly expressed in CD8+ T-cells and NK T-cells of the severe COVID-19 group. As we found that GLUT3 was associated with M01 and M03 clusters, two dominant macrophage subtypes in severe COVID-19, we further analyzed enrichment scores for glycolysis and oxidative phosphorylation (OXPHOS) pathways to explore the characteristics of glucose utilization in severe COVID-19 (Fig. 2C). Interestingly, in most immune cells, including T-cells and macrophages, enrichment scores of glycolysis were higher in the severe COVID-19 group than in the moderate and healthy control groups. In neutrophil and macrophage clusters, scores of OXPHOS were lower in the severe group than in the moderate and healthy control groups. As [^18^F]FDG uptake is determined by expression levels of GLUTs and hexokinase, the initial enzyme of glycolysis, we assume that [^18^F]FDG positron emission tomography (PET) can be used for the detection of severe COVID-19.

**Fig. 2.**
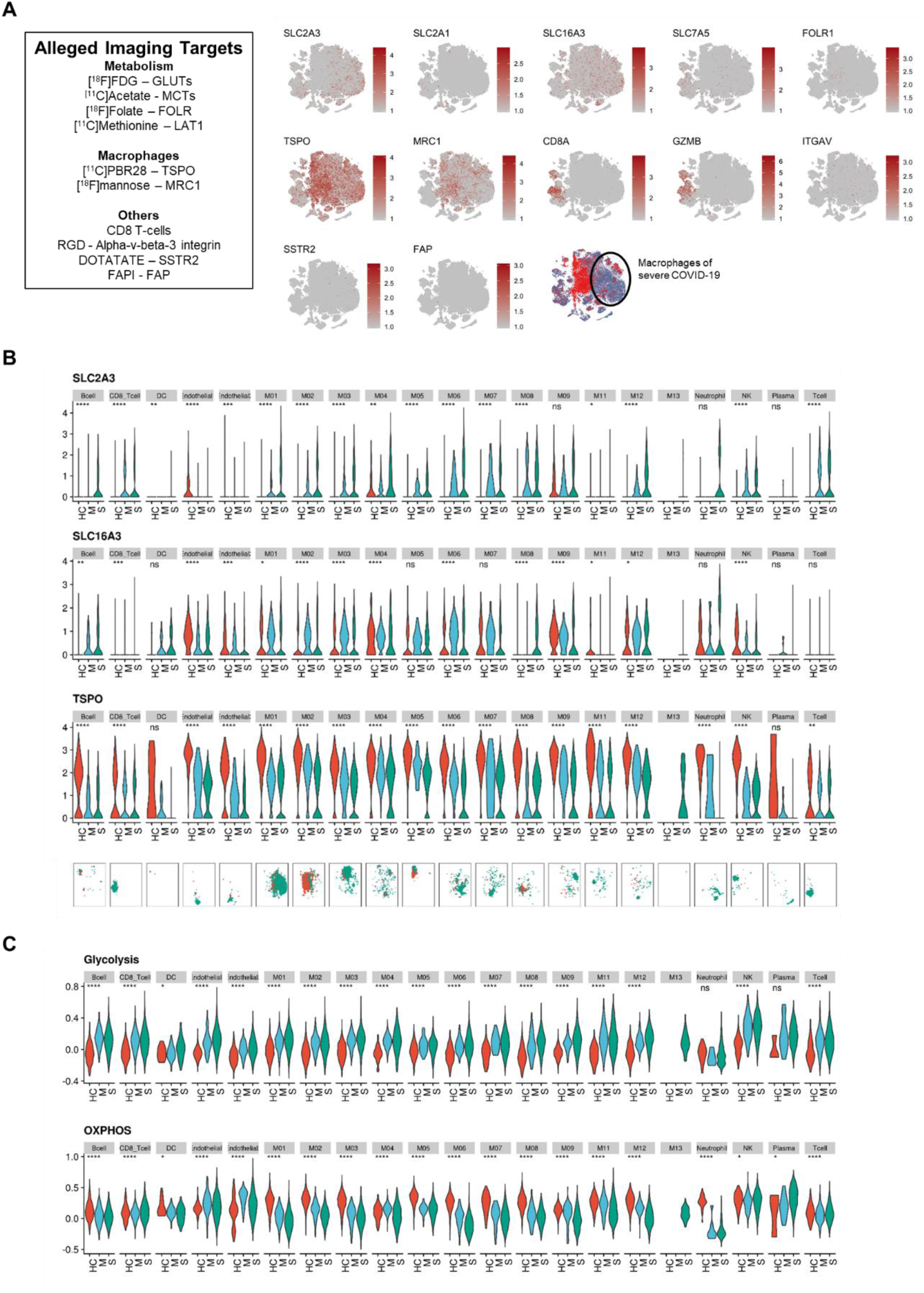
Potentially alleged imaging marker molecules of COVID-19 and metabolic pathways. **(A)** t-SNE plots showing the expression of several potentially alleged imaging targets on BAL fluid immune cells. The markers indicate the heterogeneous pattern of expression across the immune cells. The last panel indicates subtypes of macrophages abundant in severe COVID-19 patients. **(B)** Expression levels of *SLC2A3, SLC16A3*, and *TSPO* across cell clusters of three groups. (ns: p > 0.05; *: p <= 0.05; **: p <= 0.01; ***: p <= 0.001; ****: p <= 0.0001) (HC = healthy control; M = moderate COVID-19; S = severe COVID-19) t-SNE plots on the bottom panels show the distribution of each immune cell cluster. (red dot = HC; blue dot = M; green dot = S) **(C)** Enrichment scores of glycolysis and oxidative phosphorylation (OXPHOS) pathways across cell clusters of three groups.

### Exploration of ideal targets of severe COVID-19 hyperinflammation

We hypothesized that an ideal target for imaging and targeted therapy of COVID-19 would have the following characteristics: 1) highly expressed in severe COVID-19, 2) expressed on the cell surface, and 3) the existence of binding molecules. Thus, we analyzed scRNA-seq data of COVID-19 BAL fluid cells along with three different databases: the Surfaceome database (*9*), the Database of Imaging Radiolabeled Compounds (DIRAC) (*10*), and BindingDB (*11*). The Surfaceome database was constructed using publicly available data on genes to catalog all those known to (or likely to) encode cell surface proteins (www.rdm.ox.ac.uk/research/rabbitts-group); because they can be targeted by ligands or antibodies, surface proteins associated with severe COVID-19 hyperinflammation may be candidates for imaging and therapy targets. DIRAC is an open-access PET radiotracer database that provides [^18^F]-radiolabeled compounds with associated specific molecules (http://www.iphc.cnrs.fr/dirac/). BindingDB is a database of drug targets with small molecules binding to specific proteins (https://www.bindingdb.org/).

Based on the proportions of immune cells within each group (Fig. 1B), we considered M01/M03, M04, and M02 as specific macrophage subtypes for the severe COVID-19, moderate COVID-19, and healthy control groups, respectively. Venn diagrams were used to display the numbers of marker genes enriched in the specific macrophage subtypes, surface protein-encoding mRNA from the Surfaceome database, and target proteins from the PET radiotracer database (Fig. 3A). Targetable marker proteins of the M01, M02, M03, and M04 clusters were explored, and we identified specific targetable proteins included in both the Surfaceome and PET tracer databases. As a result, *SLC3A2* and *SLC2A3* in the M01 cluster and *FOLR2* in the M03 cluster were identified, indicating that [^18^F]FDG and [^18^F]-labeled folic acid derivatives may be useful for imaging severe COVID-19 (Fig. 3A).

**Fig. 3.**
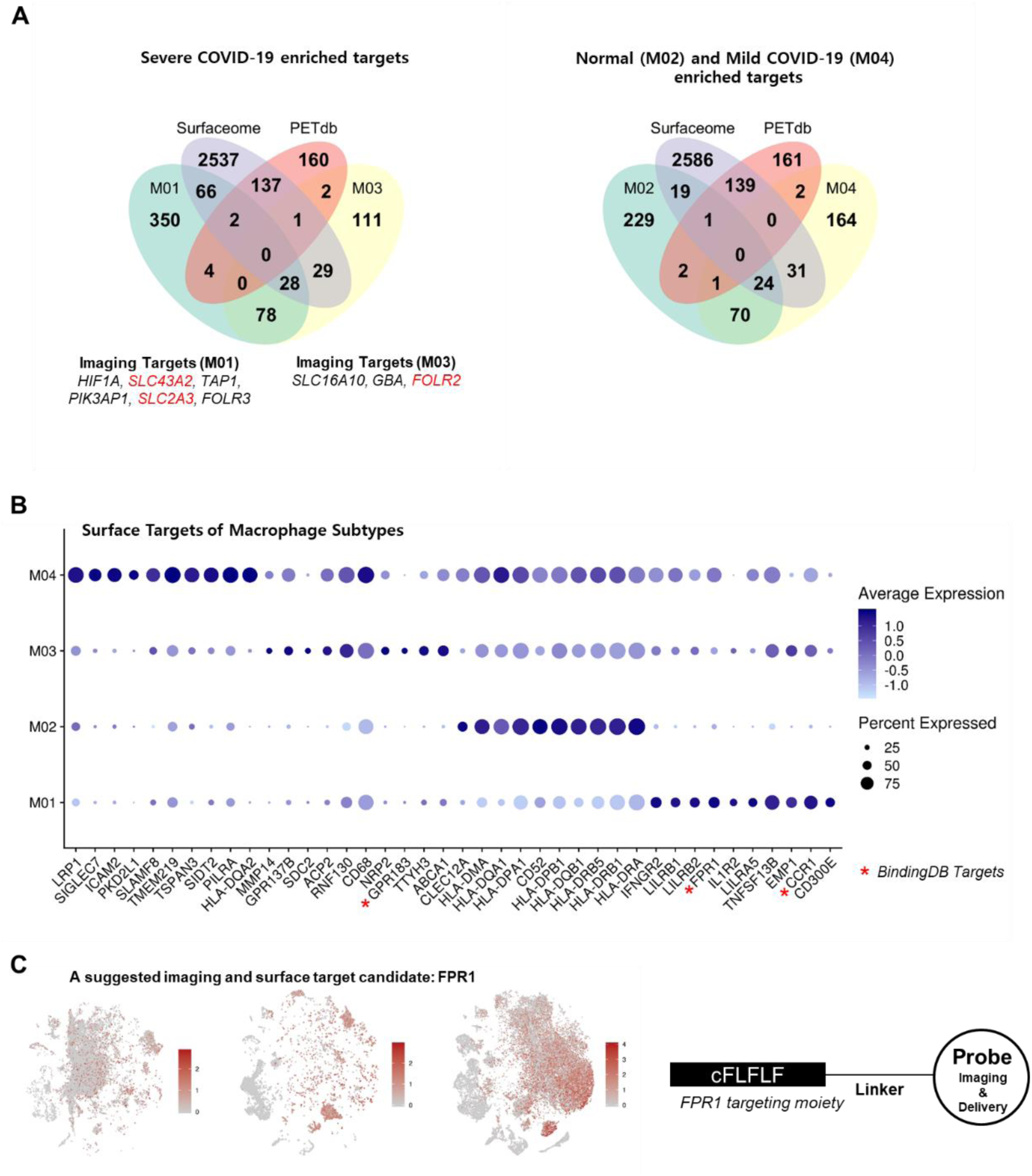
Discovery of targetable marker protein of severe COVID-19. **(A)** Venn diagrams representing intersection between markers for specific subtype of macrophages of each patient group (M01 and M03 for severe COVID-19, M04 for moderate COVID-19, and M02 for healthy control), the Surfaceome database, and the PET tracer database. The targetable surface proteins were *SLC43A2* and *SLC2A3* for M01 and *FOLR2* for M03. **(B)** A dot plot representing expression of the top 10 surface markers in specific macrophage subtypes. The dot size represents the fraction of cells expressing a specific marker in a particular cluster, and the intensity of color indicates the average expression in that cluster. **(C)** *FPR1*, which is mostly expressed in the M01 cluster, is suggested as an example of a potential imaging and surface target for severe COVID-19 patients. For the left panel, t-SNE plots represent cells acquired from healthy controls and moderate and severe COVID-19 patients, respectively.

Additionally, the top 10 surface markers of M01, M02, M03, and M04 selected by fold-changes were visualized by the average expression level and proportions of protein-expressing cells (Fig. 3B). Notably, a few surface molecules enriched in M01/M03 among the top 10 surface markers correspond to drug targets in the BindingDB, including *CCR1, FPR1*, and *GPR183. FPR1* was higher in the severe COVID-19 group than in the healthy control and moderate groups (Fig. 3C). The specific peptide ligand of *FPR1*, cinnamoyl-F-(D)L-F-(D)L-F (cFLFLF), has been used for imaging inflammation (*25*) and targeted drug delivery conjugated with nanoparticles (*26*) (Fig. 3C). Thus, we suggest that cFLFLF-based imaging or drug delivery systems can be utilized for imaging and therapy of severe COVID-19. Additionally, there are 1155 and 22 ligand candidates for *CCR1* and *GPR183*, surface targets enriched in severe COVID-19, in BindingDB, respectively. All markers of M01, M02, M03, and M04 with surface targets, PET database and targeting drug candidates are summarized in Supplementary Table 1. Accordingly, for M01/M03 subtypes, 154 surface target candidates, nine [^18^F]-labeled radiotracers, and 132 molecules associated with drug targets were identified, warranting further evaluation for utilization in imaging and therapy of severe COVID-19.

## Discussion

COVID-19 is a novel viral infection disease of which the most representative manifestation is severe pulmonary inflammation (*4*). An excessive inflammatory process is known as a main factor that leads to pulmonary destruction and even death (*27*). Many approaches to treating patients with severe COVID-19 by controlling the immune response using immunomodulators such as dexamethasone, interleukin-6 inhibitors or tumor necrosis factor blockers have been reported (*28, 29*). Currently, COVID-19 patients can be classified into mild, moderate, severe/critical diseases based on clinical symptoms according to the National Health Commission of China guidelines (*30*), and the classification of disease severity in COVID-19 is critical for the grading treatment of patients. Indeed, evaluation of the severity of the immune response is needed to select patients who urgently need anti-inflammatory treatment if the immune modulation strategy becomes an important treatment option for severe COVID-19. Moreover, it might be useful in optimizing the allocation of medical resources and preventing the occurrence of overtreatment and undertreatment in the outbreak of an epidemic. However, to the best of our knowledge, efficient indicators for the severity of COVID-19, therapeutic response, and outcome have not been fully elucidated. Some previous studies have proposed clinical symptoms, laboratory, and radiologic findings as diagnostic tools for the classification of COVID-19 severity (*31*). Our study suggests targetable molecules reflecting the different compositions of immune cells according to the severity of COVID-19, indicating that imaging and therapeutic targeting of specific molecules of hyperinflammation in severe COVID-19 may provide a new feasible strategy of stratification and precision immune modulation in COVID-19.

We explored expression of alleged imaging targets to assess the feasibility of existing molecular imaging probes for inflammation. Notably, *SLC2A3* (GLUT3) was highly expressed in macrophages enriched in severe COVID-19 patients. In addition, these macrophages showed enhanced glycolysis and relatively low OXPHOS. These findings support the evaluation of glucose metabolism in inflammatory lesions and can provide information on the immunologic response associated with severe COVID-19. Notably, by reflecting enhanced glucose metabolism in inflammatory cells, particularly macrophages, [^18^F]FDG PET has been commonly employed in clinical settings for identifying inflammatory or infection foci and evaluating the severity of inflammation (*32*). Arecent case series also showed increased FDG uptake in COVID-19-related inflammation in the lungs and lymph nodes (*33*). Our results suggest that [^18^F]FDG PET can be utilized to stratify COVID-19 patients with severe pulmonary inflammation, as enhanced glucose uptake and glycolysis were found to be associated with subtypes of macrophages in severe COVID-19 compared with moderate disease. Imaging-based characterization has advantages in reflecting the metabolic aspects of macrophages related to severe COVID-19 and evaluating the whole body to localize hyperinflammation. In addition to the pulmonary system, other organs, such as the liver or kidney, can be affected by SARS-CoV-2 (*3*). Considering the availability of [^18^F]FDG PET imaging and that can cover the whole body, it can be used for the diagnosis of systemic inflammation caused by SARS-CoV-2. Although further clinical validation is required, the degree of FDG uptake in inflammatory lesions in the lung can be used as a predictable finding of hyperinflammation by suggesting macrophage subtypes with enhanced glucose metabolism.

We further explored candidate molecules for imaging and druggable targets using databases. It is notable that *SLC2A3* (GLUT3) was selected in this analysis, and this result indicates that it is a good candidate for targeted imaging reflecting immune cells in severe COVID-19 patients. Other surface molecules associated with druggable targets were also identified; *FOLR2* was another candidate imaging target for the severe COVID-19 group. The folate receptor is a molecular target used to diagnose inflammatory diseases (*34*). This result is consistent with a previous animal study that showed the possibility of imaging *FOLR2*-positive macrophages in acute lung inflammation (*35*). In addition, we focused on surface targets that bind to druggable molecules. *CCR1* and *FPR1* were highly expressed in macrophages enriched in the severe COVID-19 group. Specific druggable molecules for these targets are indicated in BindingDB. In particular, *FPR1* is a G protein-coupled receptor expressed in macrophages. There is a previous study reporting the feasibility of *FPR1* targeted imaging to diagnose macrophage infiltration in inflammatory diseases (*36*). As FPR1 is targeted by small peptides (cFLFLF), it can be applied to active targeting drug delivery systems as well as molecular imaging (*26*). Furthermore, *FPR1* is reported to have a role in regulating or modulating the immune response in cancer and inflammation (*37*). As immunomodulation or immunoregulation has recently been emphasized as a treatment strategy in COVID-19 (*38*), *FPR1* may be an appropriate target not only for assessing disease severity but also for delivering therapeutic drugs. We identified various druggable molecules specifically binding to markers of macrophages enriched in severe COVID-19 (Supplementary Table), and the findings may be used to develop an appropriate drug delivery and imaging platform for precise immune modulation strategies in COVID-19. In other words, the present results indicate that the target molecules explored can be applied to diagnose severe pulmonary inflammation due to SARS-CoV-2 shortly.

There is a limitation in the present study. Because we analyzed cells from BAL fluid, the results may not be entirely consistent with the immune cell composition or protein expression of the lung tissue itself. Nonetheless, the characteristics or composition of cells in BAL fluid reflect immune cells of the lung associated with inflammation. Thus, targeted imaging of the identified molecules can be applied to in vivo settings. Further study is warranted to validate the feasibility of targeted theranostics of these molecules.

Taken together, we demonstrate different compositions of immune cells in BAL fluid from healthy controls and COVID-19 patients. The subpopulations of macrophages differed among the three groups. Regarding alleged imaging markers, *SLC2A3* was abundant in macrophage subtypes enriched in severe COVID-19 patients, and we identified *SLC3A2, SLC2A3*, and *FOLR2* as candidate molecules as imaging targets. In addition, various molecular targets, including *CCR1, FPR1*, and *GPR183*, are suggested as candidates for drug delivery systems as well as imaging. This work provides a resource to develop targeted imaging and therapeutic strategies for severe pulmonary hyperinflammation related to COVID-19.

## Materials and Methods

### Preprocessing scRNA-seq data

scRNA-seq data for BAL fluid were downloaded from the Gene Expression Omnibus database (GSE145926). The scRNA-seq data were scaled by log-normalization after the read counts were divided by the total number of transcripts and multiplied by 10,000. Highly variable 2,000 genes were selected using the *FindVariableFeatures* function of the Seurat package (version 3.0) (*39*). Data were then scaled to z-scores with regression of total cellular read counts and mitochondrial read counts. Cell types were determined by the graph-based clustering approach implemented by the *FindClusters* function of the Seurat package. Before clustering, dimension reduction was performed by principal component analysis, and 50 dimensions were used for clustering. The conservative resolution was set to 0.5.

### Clustering cells into each immune cell type

The *FindAllMarkers* function of the Seurat package was used to identify marker genes of the clusters, and high-ranked marker genes according to the fold-change were identified for each cluster. For data visualization, the scRNA-seq data were embedded into two-dimensional projection, t-stochastic neighborhood embedding (t-SNE). The expression levels of known marker genes, *CD68, FCGR3B, CD1C, LILRA4, KLRD1, CD3D, CD8A, MS4A1, IGHG4*, and *TPPP3* (Supplementary Figure 1), were assessed to identify cell types, and each cluster was classified into eight cell types based on the expression level: T-cell, B-cell, plasma cell, NK T-cell, endothelial cell, neutrophil, dendritic cell, and macrophage.

### Differentially expressed genes and gene set enrichment analyses

To identify marker genes, differential expression analysis was performed using the function *FindAllMarkers* of the Seurat package with the Wilcoxon rank sum test. Differentially expressed genes that were expressed in at least 10% of cells within the cluster and with a fold change greater than 0.25 (log scale) were considered marker genes. t-SNE plots and violin plots were generated using Seurat. We used Reactome to select genes of glycolysis and OXPHOS pathways to examine the overall activity of glycolysis and OXPHOS pathways (*40*). The metabolic enrichment scores of each cell were estimated by the *AddModuleScore* function of the curated gene sets of both pathways to define the metabolic profiles of each sample. Expression of marker genes and enrichment scores of metabolic pathways were compared using a Kruskal-Wallis test. For each cluster of cells, feature scores of severe, moderate COVID-19 and healthy controls were compared.

### Exploration of target molecules in open access databases

We used three different databases to discover candidate target molecules: the Surfaceome database (*9*), the Database of Imaging Radiolabeled Compounds (DIRAC) (*10*), and BindingDB (*11*). Marker genes included in the three databases were selected. Surface molecules were curated by a database downloaded from the in silico human surfaceome website (https://wlab.ethz.ch/surfaceome/). Among markers of each cell cluster, surface markers were selected by Surfaceome, and the top 10 surface targets according to the fold change were selected. The surface markers between different clusters were then compared. The DIRAC database curates [^18^F]-radiolabeled tracers and their associated molecular targets. All available image-able targets were selected and compared with markers discovered by scRNA-seq. We also used BindingDB, which provides available druggable molecules with target proteins. All statistical analyses were performed using the R program (v 3.6.1).

## Acknowledgments

None

## Author contributions

H.C., K.J.N. designed the study. H.L., K.J.N., and H.C. performed data analysis. H.J.I. interpreted and critically reviewed results. H.L. wrote the first draft. All authors wrote and revised the manuscript. All authors read and approved the final manuscript.

## Data and materials availability

The single-cell RNA-sequencing data can be downloaded from the Gene Expression Omnibus database (https://www.ncbi.nlm.nih.gov/geo/). The software and resources used for the analyses are described in the Materials and Methods.

