## Supplementary Figures for "Discovery of potential imaging and therapeutic targets for severe inflammation in COVID-19 patients"

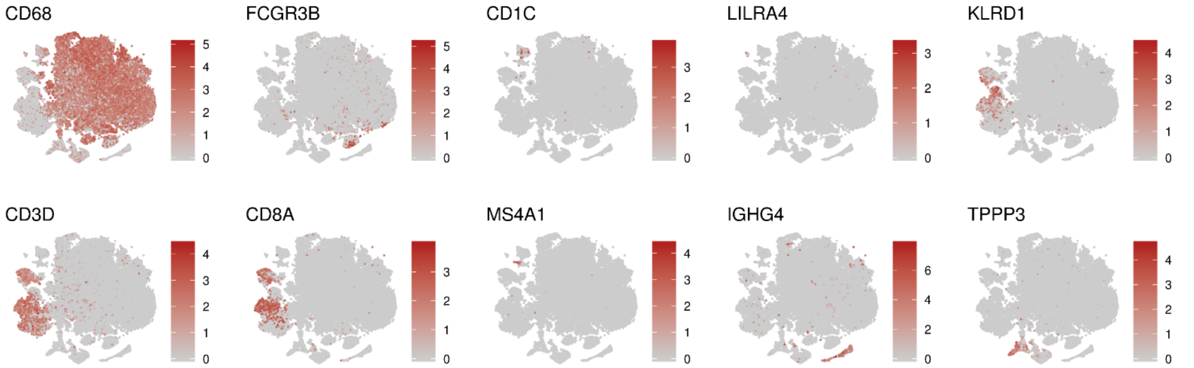


**Supplementary Fig 1. The markers for each immune cell type within BAL fluid**

t-SNE plots showing the expression of several markers on BAL fluid immune cells; *CD68, FCGR3B, CD1C, LILRA4, KLRD1, CD3D, CD8A, MS4A1, IGHG4*, and *TPPP3*.

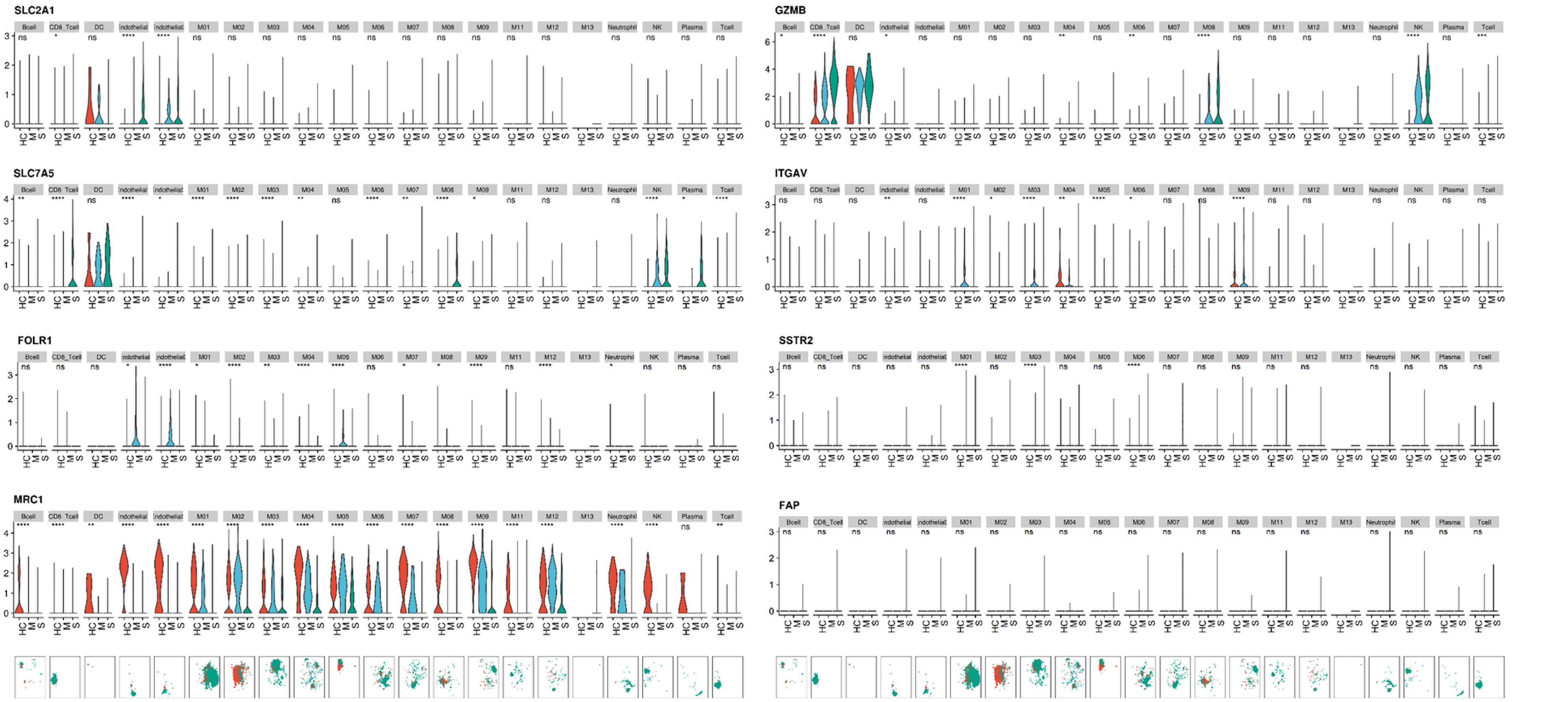


**Supplementary Fig 2. The expression of alleged imaging markers for inflammation**

The expression levels of alleged imaging markers were evaluated across immune cell clusters and compared between three groups.; *SLC2A1, SLC7A5, FOLR1, MRC1, GZMB, ITGAV, SSTR2*, and *FAP*. (ns: p > 0.05; *: p <= 0.05; **: p <= 0.01; ***: p <= 0.001; ****: p <= 0.0001) (HC = healthy control; M = moderate COVID-19; S = severe COVID-19) t-SNE plots on the bottom panels show distribution of each immune cell cluster. (red dot = HC; blue dot = M; green dot = S)
